# Epidemiological impact of hepatitis B vaccination (Monastir_ Tunisia ; 2000-17)

**DOI:** 10.1101/385880

**Authors:** Wafa Dhouib, Meriem Kacem, Grira Samia, Issam Maalel, Imen Zemni, Hela Abroug, Manel Ben fredj, Arwa Ben Salah, Souhir Chelly, Essia Green, Amira Djobbi, Asma Sriha Belguith

## Abstract

Background: In 2016, the first global health sector strategy on viral hepatitis was endorsed with the goal of eliminating viral hepatitis as a public health threat by 2030. In Tunisia, effective vaccines for hepatitis B (HBV) have been added to the expanded programme of immunization (EPI) since July 1995 for new borns. We expected to have a decreasing trend in the prevalence rate of reported HBV. Our study aimed to address the epidemiological profile of HBV, to assess trends by age and gender in Monastir governorate over a period of 18 years according to immunization status and to estimate the burden (years lived with disability YLDs) of this pathology. Methods: We performed a descriptive cross sectional study of declared HBV from January 1, 2000 to December 31, 2017 defined as having positive serologic markers for HBs Ag. All declared patients were residents of Monastir’s Governorate. EPI included two periods, the first between 1995 and 2006 following a three-dose schedule (3, 4, 9 months). The second PI cohort after 2006 following a three-dose schedule (0, 2, 6 months). Results: During 18 years, 1526 cases of HBV were declared in Monastir with a mean of 85 cases per year. We estimated a mean of 1699 declared cases per year of HBV in Tunisia. CPR was 16.85/100,000 inh being the higher in age group of 20-39 years and in men .ASR was 15.99/100,000 inh, being 35.5 in men and 8.69 in women. During the study period, declared cases among presumed immunized (PI) person against HBV were 32(2.0%). Among PI cases, 29 were from the first period and 3 were from the second. We established a negative trend over 18 years of hepatitis B. The age group of 20 to 39 was the most common with a sharply decline. Presumed not immunized (PNI) HBV cases are decreasing by years with a prediction of 35 cases in 2024. Reported HBV contributed to 1.26 YLDs per 100,000 inh. The highest rate of YLDs occurred at the age 20-39 (2.73 YLDs per 100,000 inh). During 18 years, YLDs were 114.45 in Monastir with a mean of 2293.65 YLDs of HBV in Tunisia. Conclusion, this study showed a law prevalence rate and a decreasing trend of HBV during 18 years showing an efficacy of immunization and confirming that the universal hepatitis B vaccination in Tunisia has resulted in progress towards the prevention and control of hepatitis B infection. These findings should be demonstrated in other Tunisian regions with a standardized serological profile.

## Background

Hepatitis B virus (HBV) account for more than 90% of viral hepatitis-related deaths and disability, and are the seventh leading cause of mortality globally, responsible for 1.34 million deaths in 2015(1,2). Therefore, HBV infection with its heavy burden constitutes a public health threat and is a candidate for prevention, early diagnosis, and treatment (3). In 1991, the world health organization (WHO)recommended universal infant immunization against HBV with a target for worldwide implementation by 1997. In 2016, The first global health sector strategy on viral hepatitis was endorsed with the goal of eliminating viral hepatitis as a public health threat by 2030(4). Worldwide, in 2015, the estimated prevalence of HBV infection in the age under 5 was about 1.3%(1). In Tunisia, effective vaccines for HBV have been available and added to the Expanded Programme of Immunization (EPI) since July 1995 for new borns. Since 2006, the first administration was advanced to birth following a three-dose schedule (0, 2, 6 months) instead of (3,4,9 months). since 2014, all new born received four-dose schedule(0-2-3-6 months)(5). In addition, all pregnant women are screened for HBsAg and administration of post-exposure prophylaxis is indicated for infants born to HBsAg-positive women within24 hours of birth. During prenuptial visit, seronegative couples are advised to vaccinate while it is mandatory if one is positive. Considering the high vaccination coverage in Tunisia (94%), almost all Tunisians citizens born after 1995 have been vaccinated(6).Thus, we were expecting to have a decreasing trend in the prevalence rate of reported HBV more precisely among vaccinated cohort. The evaluation of the prevalence and distribution of HBV is estimated through an epidemiological surveillance of mandatory declaration disease (7). Accordingly, this study aimed to address the epidemiological profile of HBV, to assess trends by age and gender in Monastir governorate over a period of 18 years according to immunization status (2000–2017) and to estimate the burden (years lived with disability) of this pathology.

## Methods

### Study design

We performed a descriptive cross sectional study of declared HBV during 18 years.

### Study population

All medical patients, residents of Monastir Governorate, diagnosed HBV were declared to the regional direction of primary health care from January 1, 2000 to December 31, 2017. Patients living in other governorate were not included.

### Study area

The governorate of Monastir is situated in the coastal region of Tunisia; it covers an area of 97 465 hectares and represented 0.7% of the total Tunisian territory. In 2014, the general population of Monastir governorate was 548 828 inhabitants, or 4.99% of the Tunisian population. (8).

### Data collection method and procedure

According to national guideline hepatitis management, declaration of HBV infections in Tunisia is mandatory (9). Therefore, all public or private health physicians, general practitioners or specialists, all public and private laboratories, blood donation unit, should report all diagnosed cases of HBV infections. The discovery circumstance of HBV was either clinical ( as ictere, gastroenteritis, fever..) or serological when taking blood for prenuptial examination, pregnancy or others .All patients with serologic markers positive for HBsAg were recorded in a database in the regional direction of primary health. This database contained patient demographic characteristics (age, sex, residence).

The first presumed immunized(PI) cohort was defined as persons who were born and vaccinated since the starting of EPI (1995) to 2006 and following a three-dose schedule (3,4,9 months). The second PI cohort was defined as persons who were born and vaccinated after 2006 following a three-dose schedule (0, 2, 6 months).

In this study, we considered that the risk of underreporting was constant since the reporting physicians during this period were almost the same.

The burden of hepatitis is measured in Disability Adjusted Life Years (DALYs) which is the sum of YLDs (years lived with disability) and YLLs (years of life lost).

The estimation of YLD for a given disorder is a product of epidemiological data that accommodates the number of people affected as well as the severity and disability associated with their symptoms. That is, YLDs are calculated by multiplying the prevalence of a disorder by its severity and comorbidity-adjusted disability weight.

YLDs = Prevalent cases × Disability weight

Disability weights quantify the severity of a health state on a scale of zero (complete health or no disability) to one (complete disability, equivalent to death). Disability weights for GBD 2013 were derived from a pooled analysis of data from the GBD 2010 Disability Weights Measurement Study(10).

### Statistical analysis

Data were verified and analyzed using IBM SPSS Statistics version 22.0 software (IBM Corp., Armonk, NY, USA). Categorical variables were reported as numbers and percentages. The Crude Prevalence Rate (CPR) was calculated based on Tunisian National Institute of Statistics data and was expressed as the number per 100,000 inhabitants(inh)(11)The Age-Standardized prevalence Rate (ASR) per 100,000inh was calculated using the world standard population according to the World Health Organization statement of 2013(12). The average population was calculated as follows: (Monastir population in 2004 + Monastir population in 2014)/2. Linear regression was used to calculate the slope ‘b’ of the least-squares line for estimating the trends in notified disease according to sex and age group. Predictions were performed using the Age Period Cohort based on Poisson regression.

A p-value of <0.05 was considered statistically significant.

### Ethical considerations

The study was conducted under Good Clinical Practice conditions and according to ethical standards collections. Each patient in the study was assigned a unique identifying code and all documents were labelled accordingly to maintain anonymity.

Case investigations around a case are conducted by health workers responsible for this task to diagnose and manage cases of hepatitis according to ethical standards.

## Results

During 18 years, 1526 cases of HBV were declared in Monastir with a mean of 85 cases per year. We estimated a mean of 1699 declared cases per year of HBV in Tunisia. CPR was 16.85/100,000 inh being the higher in age group of 20-39 years and in men. ASR was 15.99/100,000 inh, being 35.5 in men and 8.69 in women. During the study period, declared cases among PI person against HBV were 32(2.0%) with one case in 2004 and 3 in 2014 (Table 1). Among PI cases, 29 were from the first PI cohort and 3 were from the second PI cohort. Figure1 showed that presumed not immunized (PNI) HBV cases are decreasing by years with a prediction of 35 cases in 2024 with an increasing number of PI cases especially between 2014- 2017 with a prediction of 12 cases in 2024. We established a negative trend over 18 years of HBV (b= −3,977; r = −0.804; p<10^−3^). The age group of 20 to 39 was the most common with a sharply decline. (b= -3,768; r = - 0.907; p<10^3^) (figure2).

**Table 1:** Crude and age standardized prevalence rates, by age groups, gender and immunization status of declared HBV (2000-2017)

|  | **Cases(%)** | **CPR \*** | **ASR\*** |
| --- | --- | --- | --- |
| **Over all** | 1526(100) | 16.85 | 15.99 |
| **Age groups (Years)** |  |  |  |
| 0-19 | 109(7.1) | 3.24 |  |
| 20-39 | 1121(73.5) | 36.40 |  |
| 40-59 | 278(18.2) | 15.22 |  |
| ≥ 60 | 18(1.2) | 2.30 |  |
| **Gender** |  |  |  |
| Men | 1213(79.5) | 26 .66 | 35.5 |
| Women | 313(20.5) | 6.95 | 8.69 |
| **Immunization status\*\*** |  |  |  |
| Immunized | 32(2.0) |  |  |
| Not immunized | 1494(97.9) |  |  |
| **At 2004 <sup>a</sup>** |  |  |  |
| Immunized (aged ≤9 years) | 1 | 1.17 |  |
| Not Immunized (aged > 9 years) | 117 | 31,51 |  |
| **At 2014<sup>b</sup>** |  |  |  |
| Immunized (aged ≤19 years) | 3 | 15.49 |  |
| Not Immunized (aged > 19 years) | 45 | 12.66 |  |

*\*: per 100000 inhabitant/years; **CPR**: crud prevalence rate; **ASR**: age standardized rate;*
*\*\* :immunized :cases born after 1995. /not immunized :cases born before 1995.*
(a):after 9 years of starting the national vaccination programs(1995)
(b) after19 years of starting the national vaccination programs(1995)

**Figure 1:**
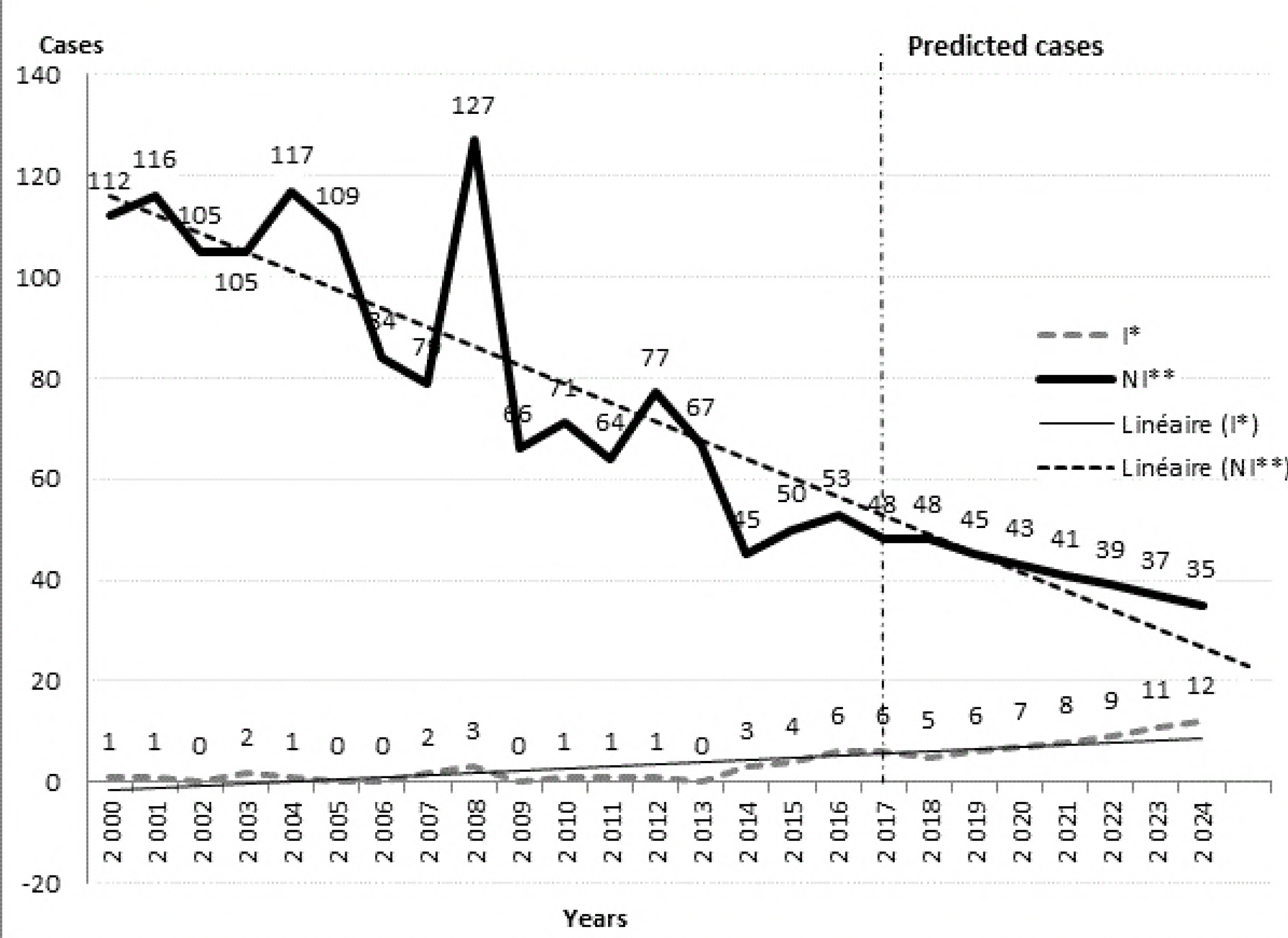
Distribution of immunized and not immunized HBV cases by year(2000-2017) with prediction at 2024. ^*^:immunized: *cases born after 1995;* ^**^*:not immunized :cases born before 1995*.

The relative risk of expressing clinical disease among women was 7.68 than in men (figure3). Clinical cases were 185 with 1341 of serological cases. Reported HBV contributed to 1.26 YLDs per 100,000 inh. The highest rate of YLDs occurred at the age 20-39 (2.73 YLDs per 100,000 inh). Men had a greater YLDs with 1.99/100,000 inh than women 0.52/100,000 inh. During 18 years, YLDs were 114.45 in Monastir with a mean of 2293.65 YLDs of HBV in Tunisia.

**Figure 3:**
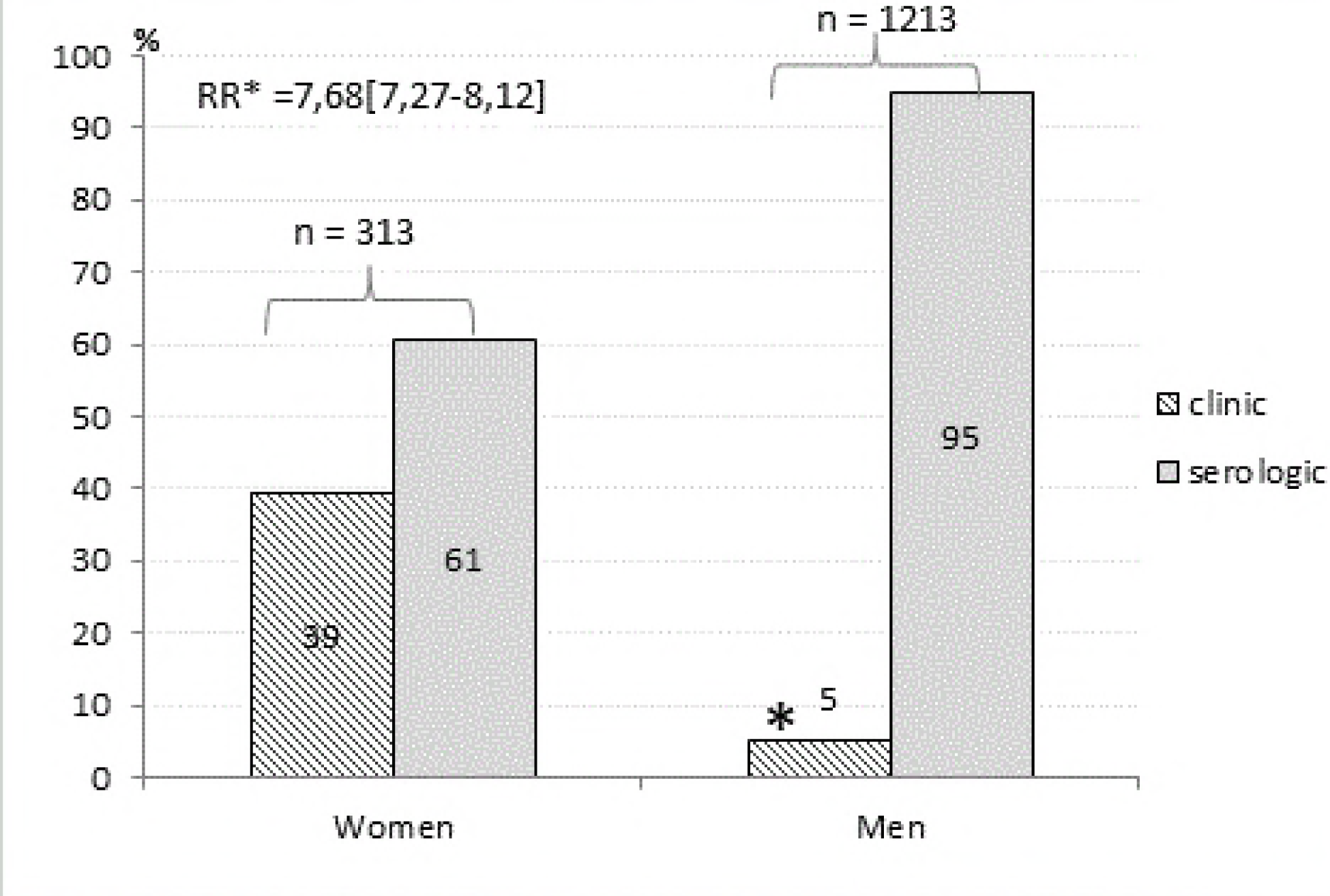
Distribution of clinical and serological declared HBV cases among men and wom en during 18 years in Monastir.

## Discussion

This is the first epidemiological survey of HBV infection conducted in Tunisia after the implementation of the EPI in 1995.Results of this study contribute to evaluation of the impact of vaccination, providing epidemiological data of HBV infection and estimating the burden of this disease during 18 years .The main results indicated that the overall prevalence rate of HBV infection is decreasing (r = -0.804) and that the major distribution of HBV was in 20-39 years group with a masculine predominance. The PNI HBV cases are decreasing by years with an increasing number of PI cases.

CPR was 16.85/100,000 inh with an ASR of 15.99/100,000 inh which is concordant to the national level. In fact, Tunisia is located in the Eastern Mediterranean Region, known as an area with an intermediate prevalence of HBV infection in 2009 (13) and low prevalence in 2015 (14).

The low prevalence rate of HBV infection in less than 20 years of age is attributed to vaccination against HBV infection and immunity. In this study, the major distribution and the upper decline of HBV was in 20-39 years group. The high rate of viral hepatitis prevalence in this age group can be explained either by the fact that it is the age of screening or by the fact that the majority did not benefit from vaccination. An interesting Tunisian study finding conducted in Sousse showed a linear decline of sero-protection rates with increasing age, from 79⋅3% to 52⋅4%, respectively, 12 and 17 years after primary vaccination(15). In addition, this age group is the commonly exposed to risk factors like unprepared sexual activity, transfusions, and history of surgery or dental work. The upper decline of HBV was in 20-39 years group. Same conclusion are shown over two time periods considered (1957-1989) and (1990-2013) in the eastern Mediterranean regions (16). Unlike the results of this study, findings of 2 similar studies in Iran (MENA) showed an upward trend of HBV infection (17, 18). One reason of this contradiction may be due to the differences in the years of two studies and the efficacy of the preventive system. A study conducted in our same governorate of Monastir (2002-2013) showed that the trend of viral hepatitis hospitalized in the university hospital of Monastir has significantly increased (19). This difference in trend can be explained either by an under-reporting or under detection of viral hepatitis or by increased trend of severe viral hepatitis requiring hospitalization against a general decline in viral hepatitis prevalence.

There was a tremendous difference in the prevalence of the disease between males and females which was consistent with a study done at the University Hospital of Monastir (2002-2013)(19). This might be related to the fact that men are more exposed to risk factors such as transfusions (road and work accident), sexual behavior and imprisonment (20). However; we have to mention that women attend more screening than men due to cesarean surgery, the tests done during pregnancy. Another important contributing factor responsible for a higher prevalence among the male population of the present study is that plasma disappearance rate for HBsAg in males is lower than females who have a better immune response (21–23).Other studies showed the same distribution such in Northern India (24) and France(25).

The PNI cases are decreasing by years with an increasing number of PI cases. In light of these results, we believe that the decrease found in PNI persons is connected to the decrease in HBV carriers in the area since introduction of universal vaccination, which therefore reduces the risk of contagion chain for those not immunized by reducing the spread of HBV in the area.

On the other hand, our results showed that the PI persons are increasing with a percentage of 90% of them coming from the first cohort Immunization .These results lead to evaluate the vaccination status. Between 2000-2017, YLDs were 114.45 in Monastir with a mean of 2293.65 YLDs of HBV in Tunisia .Between 1990 and 2013, it was 874(612-1181) thousands YLDs in north Africa and middle east according to The global burden of viral hepatitis (2). This difference might be due to the difference of time and duration of period studies or might be explained by the decreasing trend of HBV in Tunisia.

**The limitation** of this study concerns the collected data based on passive surveillance system, therefore underreporting and lack of completeness are inevitable in this study. In addition, thi study was limited in the gouvernorate of Monastir which could not reflect the epidemiological national status. Confirmatory tests have not been used for definitive diagnosis of subjects including recheck of HBsAg for chronic HBV infection diagnosis, thus some reported cases may be false positive or occult HVB(26).In the monitoring system, specifying the number of received vaccine doses were not mentionned.

Through this study, we encourage to Take more seriously the vaccination against HBV infection for high-risk groups, other family members of positive cases and immunization in the prenuptial visits .In addition, Physicians, health personnel and laboratories must be aware of the obligation to comply with the declaration of hepatitis and all other reported diseases.

In conclusion, this study showed a law prevalence rate and a decreasing trend of HBV during 18 years showing an efficacy of immunization and confirming that the universal hepatitis B vaccination in Tunisia has resulted in progress towards the prevention and control of hepatitis B infection. These findings should be demonstrated in other Tunisian regions with standardized serological profile.

## Acknowledgments

The authors thank the following people for their help in data collection and providing them: the team of the regional direction of primary health, department of Epidemiology and Preventive Medicine at the university of Monastir, department of family medicine; Faculty of medicine of Monastir, Tunisia. This study received no financial support. We offer this paper to our eminent Professor Soltani Mohammed.

## Ethics approval and consent to participate

The study was conducted under the approving of the regional direction of primary health according to ethical standards collections with maintaining anonymity of each patient.

## Competing interests

The authors declare that they have no competing interests.

